# A Novel Amino Acid Deletion and Substitution in *amrB* Gene Associated with Gentamicin Susceptibility in *Burkholderia pseudomallei* from Malaysian Borneo

**DOI:** 10.1101/2023.05.31.543095

**Authors:** Ainulkhir Hussin, Sheila Nathan, Muhammad Ashraf Shahidan, Mohd Yusof Nor Rahim, Mohamad Yusof Zainun, Nurul Aiman Nafisah Khairuddinb, Nazlina Ibrahim

## Abstract

*Burkholderia pseudomallei* is a highly pathogenic saprophyte that is intrinsically resistant to a wide variety of antibiotics. Resistance to gentamicin is considered as an earmark of *B. pseudomallei.* However, rare susceptible strains have been isolated in certain regions due to gene mutations. Currently, data on the susceptible strains’ prevalence and the actual causal mutations are still scarce, particularly in Malaysian Borneo. A pool of *B. pseudomallei* isolates (*n* = 46) were screened for gentamicin susceptibility and phenotypically confirmed using the gradient minimum inhibitory concentration method. Three isolates were gentamicin-susceptible strains and were identified as having originated from Bintulu, Sarawak, Malaysian Borneo. The a*mrB* gene mutation in these mutant strains was analysed, and the effect of amino acid substitution on the stability of the amrB protein was determined by using *in silico* analysis. The mutagenesis analysis identified a polymorphism-associated mutation, g.1056T>G, and two susceptible-associated mutations identified as novel in-frame amino acid deletion p.Val412del and amino acid substitution p.Thr368Arg that compromised gentamicin resistance. *In silico* analysis using amrB homology-modelled and AlphaFold-solved structures proposed the role of p.Thr368Arg amino acid substitution in conferring GEN susceptibility by other mechanisms than destabilising the structure of amrB protein, which is most probably due to the mutation’s location in the highly conserved region. The findings have shed light on the phenotypic characteristics and mutations involved in the *amrB* gene of the gentamicin-susceptible *B. pseudomallei*.

## INTRODUCTION

Most of the *B. pseudomallei* strains are intrinsically resistant to aminoglycosides, including gentamicin (GEN) antibiotic (1). Based on the fact that accurate and specific gold standard culture method is paramount for the an accurate treatment and effective management of infection in melioidosis patients to prevent mortality, the *B. pseudomallei* screening agars are mostly supplemented with GEN antibiotic to expedite laboratory diagnosis by preventing the growth of other gram-negative microorganisms and improve the agar’s specificity (2–5). However, there are certain regions that have been reported to isolate gentamicin-susceptible (GEN^s^) *B. pseudomallei* strains, particularly in Sarawak, Malaysia (6, 7) and North-eastern Thailand (8, 9), which may complicate the melioidosis diagnosis due to the occurrence of false-negative culture results, causing *B. pseudomallei* infection diagnosed as other bacterial infections with melioidosis-like symptoms such as tuberculosis (10–15), community-acquired pneumoniae infection (16) or *Pseudomonas sp.* infection (17).

Several studies have shown that the GEN^s^ strains’ emergence has been caused by mutations in the resistance-nodulation-division (RND) family of multidrug efflux system, the amrAB-OprA gene, which impairs the tripartite AmrAB-OprA efflux protein function to extrude aminoglycosides, macrolides and fluoroquinolones groups of antibiotic (18, 19). The amrAB-OprA efflux pump system contains two essential components, including the *amrA* and *amrB* genes (20). GEN-susceptibility can be caused by any mutation within this efflux two genes, and previously reported mutations include a complete deletion of the amrB gene sequence (19), a complete genomic deletion of AmrAB-OprA genes, a lack of AmrAB-OprA expression (9), missense mutations (7), and frameshift mutations (18) in the *amrB* gene. In addition, a mutation in the BpeAB-OprB pump (21) has been linked to GEN-susceptibility in *B. pseudomallei* KHW strain. It is however strain-specific, as Mima and Schweizer (22) discovered it to be absent in the *B. pseudomallei* 1026b strain. For that reason, most of the GEN susceptibility-causing mutation studies were focused on the AmrAB-OprA RND instead of the BpeAB-OprB RND due to the unclear role of the latter in conferring GEN susceptibility.

A complete deletion of an entire gene, a frameshift mutation, or a nonsense mutation have been shown to result in the production of non-functional proteins (23). A nonsense mutation leads to the production of a stop codon, which results in the production of a shortened protein that is most likely non-functional (National Human Genome Research Institute, https://www.genome.gov/genetics-glossary/Nonsense-Mutation), while a frame shift mutation due to insertion or deletion of nucleotide bases not in multiples of three may cause wrong protein translation or the creation of a stop codon (National Human Genome Research Institute, https://www.genome.gov/genetics-glossary/Frameshift-Mutation). In contrast, the role of missense mutations in causing unfunctional proteins may vary (24). Although experimental mutagenesis analysis using constructed mutant confirmed that several missense mutations in the *amrB* gene cause GEN-susceptibility in *B. pseudomallei* (7, 18), the mechanism by which the identified missense mutations to confer GEN susceptibility remains unknown. Using *in silico* analysis, there are several mechanisms of missense mutation that can affect protein function and alter phenotypic character, such as protein structure stability perturbance, the location of the mutation at the active site or key interaction site for binding, or in the highly conserved region (25–27). To date, most dysfunctional proteins (58% to 93%) were caused by loss of stability, which is commonly indicated by high average energy changes (ΔΔG) (>4.8 kcal/mol) (25).

Therefore, this study was designed to screen and determine the prevalence of GEN^s^ strains in a pool of *B. pseudomallei* clinical isolates from Malaysian Borneo, including Sabah and Sarawak. Next, the MICs using gradient strip method were used to confirm the phenotype of presumed GEN^s^ *B. pseudomallei* isolates that failed to grow on screening agar. The mutations in the *amrB* gene that confer the GEN^s^ phenotype were further determined. *In silico* analysis was used to calculate the ΔΔG caused by the amino acid substitution in the homology-modelled and AlphaFold solved amrB protein to determine the mechanism of identified missense mutation to confer GEN susceptibility.

## MATERIALS AND METHODS

### Ethical considerations

Ethical approval to conduct the study was obtained from the Medical Research and Ethics Committee (MREC), Ministry of Health, Malaysia (NMRR-16-1757-32197).

### Bacterial strains

All clinical *B. pseudomallei* isolates used in this study were from patients’ specimens at Queen Elizabeth Hospital, Sabah, and Bintulu Hospital, Sarawak, Malaysia, neighbouring regions of Indonesia and Brunei Darussalam (Fig S1).

### *B. pseudomallei* GEN^s^ isolates screening and selection

A total of 41 Sabah isolates and five Sarawak isolates initially confirmed as *B. pseudomallei* using the VITEK^®^ 2 analyzer and 16S rRNA sequencing were subjected to the GEN^s^ *B. pseudomallei* screening using Ashdown’s agar (ASA). Bacterial isolates that did not grow on ASA were designated as suspected GEN^s^ strains, while those that did grow were designated as suspected gentamicin-resistant (GEN^r^) strains. All suspected GEN^s^ strains, along with two suspected GEN^r^ strains each from Sabah and Sarawak, were subjected to phenotypic and genotypic characterisation. Clinical summaries from laboratory requisition forms were extracted and analysed.

### Phenotypic characterisation

Phenotypic characterisation was done by the establishment of GEN MIC against both of the GEN^r^ and GEN^s^ *B. pseudomallei* strains. MIC determination using Etest^®^ strips (bioMérieux, France) was performed according to the manufacturer’s instructions (28, 29). Briefly, plates containing Etest^®^ strips and *B. pseudomallei* strains were incubated at 37°C for 18 to 24 hours prior to MIC reading and interpretation. The interpretation value for GEN MIC was defined as ≤4 µg/mL (susceptible), 8 µg/mL (intermediate) and ≥16 µg/mL (resistant) (30, 31). The Etest^®^ MIC values between the standard two-fold dilutions suggested by the Clinical and Laboratory Standards Institute (CLSI) (31, 32) have been rounded up to the next upper two-fold value for antibiotic susceptibility test interpretation (28).

### DNA extraction and nucleotide sequencing

DNAs from pure bacterial cultures were extracted using the QIAamp DNA mini kit (Qiagen, Germany). Following that, amplification of the *amrB* gene was carried out by using high-fidelity thermostable DNA polymerase and primers as previously described (7). The standard PCR mixture contained 10 µM of each primer, 2 mM dNTP, and 20 ng/µL template. After the initial denaturation step for 2 min at 94°C, the reaction mixtures were cycled 30 times through denaturation at 98°C for 10 s, annealing at 58°C for 15 s, and extension at 68°C for 1 min.

Amplified DNA fragments were purified using the QIAquick^®^ Gel Extraction Kit (Qiagen, Germany). Then, the bacterial DNA was quantified spectrophotometrically. An amount of 150 ng/µL was used for sequencing. The sequencing process was done by Apical Scientific Sdn. Bhd. (Selangor, Malaysia). Forward and reverse DNA sequence assembly and analysis were performed using the BioEdit Sequence Alignment Editor (33, 34). The trimmed and aligned 16S rRNA and *amrB* nucleotide sequences were initially queried via BLASTN 2.9.0+ (35–37) prior to submission via BankIt to The National Center for Biotechnology Information (NCBI) nucleotide database (38, 39).

### Determination of amrB gene conserved domain

The determination of the *amrB* gene conserved domain was carried out using Conserved Domain Search (CD-Search) in Conserved Domain Databases (CDD) (CDD v.3.20-59693 PSSMs) at the National Center for Biotechnology (NCBI) website (40). The Expect Value (E-value) threshold was set at 0.01, and the maximum number of hits was set at 500 (41, 42). *B. pseudomallei* strain K96243 *amrB* gene sequence (GenBank^®^ accession no: BX571965) was used as the query sequence (43, 44). The model for conserved region determination was chosen based on the best E-value. If two or more models had similar E-values, the bit score was used as the determinant (National Center for Biotechnology Information, https://www.ncbi.nlm.nih.gov/Structure/cdd/cdd_help.shtml - <u>RPSBWhat</u>).

### Determination of mutations within *amrB* gene

Sequencing results of the GEN^s^ *B. pseudomallei amrB* gene were compared with published *amrB* gene sequences of the reference 1026b strain (GenBank^®^ accession no: AF072887) (19, 45), K96243 strain (GenBank^®^ accession no: BX571965) (43, 44), Sabah clinical GEN^r^ strains (QEH 20 and QEH 26) and the Sarawak clinical GEN^r^ strains (QEH 56 and QEH 57) to identify polymorphism-associated and susceptibility-associated mutations using BioEdit Sequence Alignment Editor Version 7.2.5 software (33, 34).

The polymorphism-associated mutation is usually associated with a synonymous mutation that codes for a similar amino acid within the *amrB* gene sequence. The mutation is commonly found in both *B. pseudomallei* GEN^r^ and GEN^s^ strains, as a result of gene heterogeneity and unlikely to confer GEN susceptibility. In contrast, a mutation that specifically found in the *amrB* gene sequences of GEN^s^ strains was referred as a susceptibility-associated mutation.

**TABLE 1.**
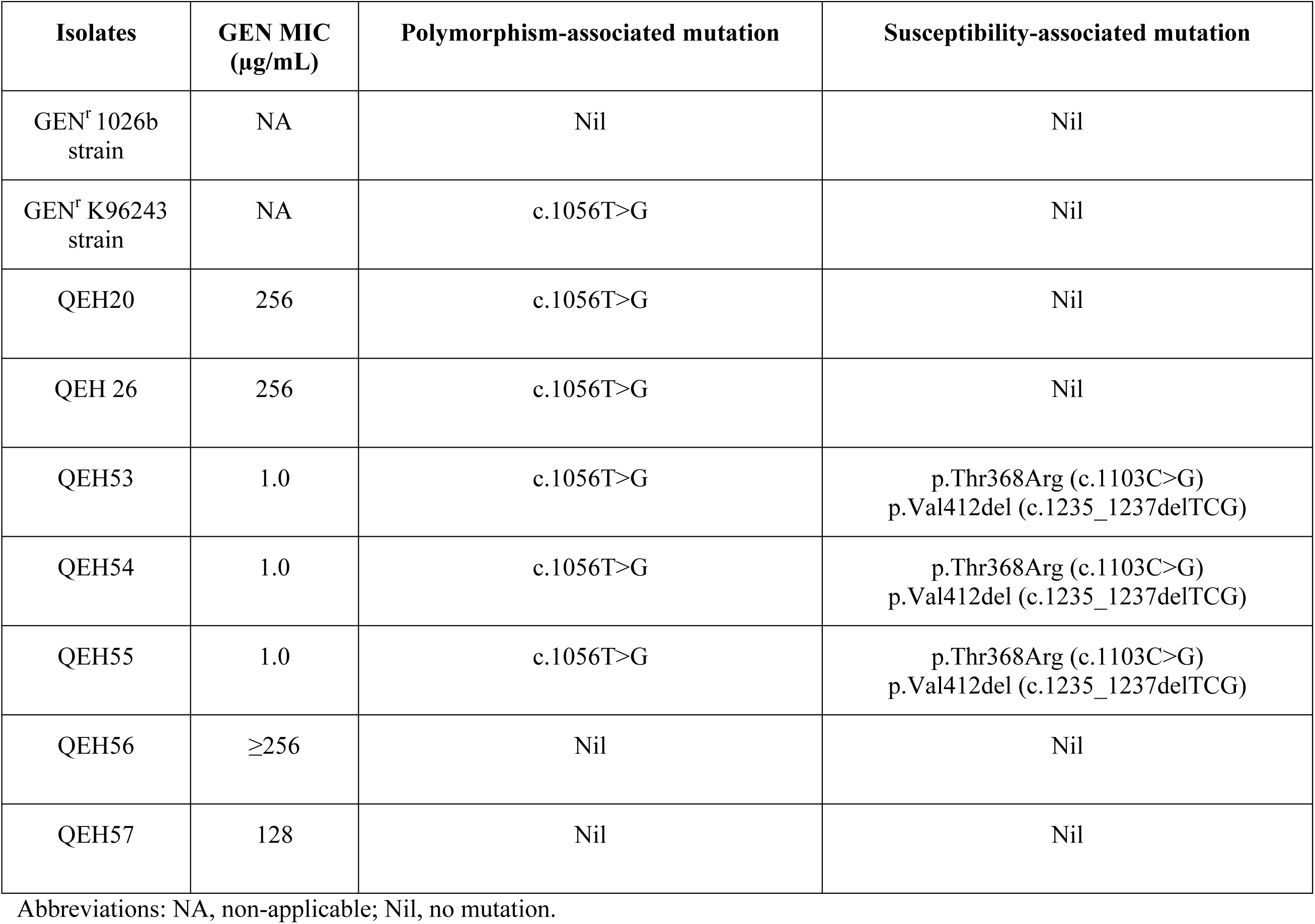
Mutations associated with polymorphisms and GEN-susceptibility in the *amrB* gene sequences

### amrB protein structure prediction by homology modelling

The complete amrB amino acid sequence for the *B. pseudomallei* K96243 strain was obtained from the GenBank^®^ database (GenBank^®^ accession no: BX571965) (43, 44), and served as the query sequence for the construction of the amrB protein using the existing template submitted to the SWISS-MODEL Template Library (SMTL) (46–50). The quality of the template model was estimated using the Global Model Quality Estimate (GMQE) (51) and Quarternary Structure Quality Estimate (QSQE) (52). The GMQE and QSQE scores are numeric values between 0 and 1, with 1 being the highest model quality (51, 52). ProMod3 was used to model proteins based on the best template (48, 53). The Qualitative Model Energy Analysis (QMEAN) scoring function (transformed to Z-score) (54) was used to determine the absolute quality of the three-dimensional (3D) model, whereas the Ramachandran plot was used to validate the structure using the MolProbity version 4.4 SWISS-MODEL programme (55). The QMEAN Z-score near zero indicates the best correlation between the model and a similar-sized experimental structure, whereas a Z-score less than -4.00 indicates the poorest model quality (SWISS-MODEL, https://swissmodel.expasy.org/docs/help). A value of more than 90% favoured amino acid residues in the Ramachandran plot indicates the good-quality structure of the built model (56, 57).

### amrB protein structure prediction by AlphaFold

The complete amrB amino acid sequence for the *B. pseudomallei* K96243 strain was retrieved from the GenBank^®^ database (GenBank^®^ accession no: BX571965) (43, 44), and served as the query sequence for amrB protein structure prediction using AlphaFold (58) via the ColabFold programme--AlphaFold2 using MMseqs2 v1.4 (59, 60).

The predicted protein structure quality was evaluated by the score of Predicted Local Distance Difference Test (pLDDT) and Predicted Aligned Error (PAE) (Å). The pLDDT score is a numeric value between 0 and 100, with 100 being the highest local structure confidence. A lower PAE value indicates more confident structure and vice versa.

### Evaluation of predicted structures by homology modelling and AlphaFold using pairwise similarity determination

The pairwise similarity between amrB protein structures predicted by using homology modelling and AlphaFold was determined by superimposing methods using TM-align version 20190822 (61, 62). The TM-score value is a numeric value between 0.0 and 1.0, with 1.0 being perfect identical structures with a similar folding rate. The TM-score achieved between 0.0 and 0.3 indicates the structure similarity was random, whereas the TM-score between 0.5 and 1.0 shows both structures were highly identical with a similar folding rate. TM-align programmes evaluated the aligned residue geometry of both structures using the root-mean-square-deviation (RMSD) value just after the superimpose function was performed. The aligned residues fit within the low RMSD value, which indicates high residue geometry with accurate structure superimposition, while the high RMSD value shows uneven residue geometry and inaccurate structure superimposition.

### ΔΔG calculation

The stability of the mutated protein structure was determined by using the mutant model building function in YASARA version 21.12.19 (63, 64). A predicted amrB protein structure model was used as the template for the mutant modelling. Energy minimisation was performed to generate a thermostable structure without van der Waals clashes using RepairPDB in FoldX (65). Structural models of the *B. pseudomallei* amrB protein mutant were done using the Mutate Single Residue protocol and energy changes were carried out using FoldX as implemented in the YASARA programme suite (65, 66). The conformer isomerism analysis was done using four conformation isomers. In this study, ΔΔG = 0 corresponds to the same stability as the wild-type protein, variants with ΔΔG value >0 correspond to those that are predicted to destabilise the protein structure, while variants with ΔΔG value <0 correspond to those that are predicted to stabilise the protein structure. A very large ΔΔG (> 10 kcal/mol) might indicate a problem in the overall structure of the model.

### Maps

ArcGIS ArcMAP version 10.2.1 (ESRI, USA) (67) was used to generate the map.

## RESULTS

### Screening, selection and phenotypic characterisation of GEN^s^ *B. pseudomallei*

The GEN antibiotic contained in ASA agar has no effect on the replication of the wild-type GEN^r^ *B. pseudomallei* strains. The isolation rate of GEN^s^ isolates was 6.52 percent, with only three Sarawak isolates from a pool of 46 that were confirmed as *B. pseudomallei* isolates being unable to grow on ASA agar and thus presumed to be GEN^s^ strains. With GEN MIC values of 1.0 g/mL, the presumed GEN^s^ isolates were confirmed as GEN^s^ strains. The presumed GEN^r^ isolates, on the other hand, had a GEN MIC range of 128 to ≥256 g/mL, confirming their resistant phenotypes (Fig. 1 & Table S1). There was no significant difference between the symptoms or risk factors of GEN^r^ and GEN^s^ infections observed (Table S1).

**Fig. 1.**
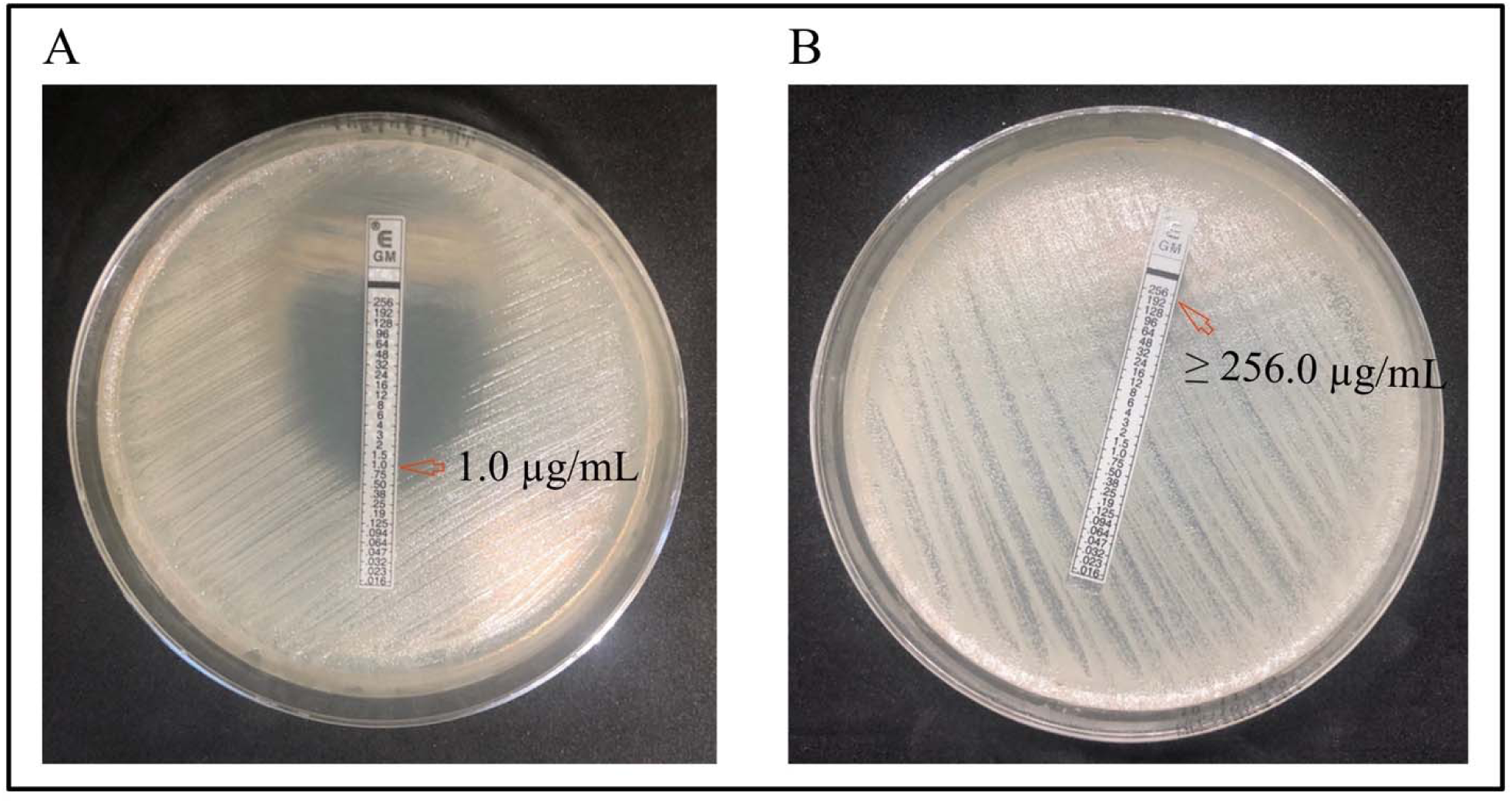
MIC values of GEN antibiotic against selected *B. pseudomallei* isolates. (A) QEH53 strain (GEN^s^) with GEN MIC value of 1.0 µg/mL) and (B) QEH56 strain (GEN^r^) with GEN MIC value of ≥ 256.0 µg/mL.

### The *amrB* gene was successfully amplified and sequenced

The partial *amrB* genes of *B. pseudomallei* isolates with an approximate size of 400 bp were successfully amplified and sequenced (Fig. S2). The accession numbers MN508906-MN508912 have been assigned to these sequences in the GenBank database (Table S2).

### The highly conserved domain in the amrB protein successfully determined

The highly conserved domain in the amrB protein was successfully identified by comparing the query amrB protein sequence to conserved domain models submitted to the CDD NCBI database. The protein sequence of amrB was strikingly similar to that of the MexY subunit of the MexXY-OprM *P. aeruginosa* aminoglycoside efflux system, which mediates resistance to aminoglycoside antibiotics such as GEN (data not shown). The majority of the amrB protein sequence is highly conserved, similar to the MexY conserved domain of *P. aeruginosa* (Fig. S3). A mutation within a highly conserved region is thought to be the primary cause of protein destabilisation, resulting in a non-functional protein.

### Polymorphism-associated mutation and susceptibility-associated mutations were successfully identified

All GEN^s^ *B. pseudomallei* strains contained a polymorphism-associated mutation in the *amrB* gene (c.1056T>G) that can be considered a genetic variation or heterogeneity that was contributed by a synonymous point mutation. The mutated codon was translated to the similar amino acid, alanine (Ala) in a less conserved region that was predicted not to confer gentamicin-susceptibility (Fig. S4 & S5) and linked to the gene polymorphism. This mutation was also found in the *amrB* gene sequences of GEN^r^ isolates, including *B. pseudomallei* from Sabah (QEH20 and QEH26) and *B. pseudomallei* strain K96243 (Table 1 and Fig. S5).

Conversely, we identified two susceptibility-associated mutations that confer the GEN^s^ phenotype, including a missense mutation p.Thr368Arg (c.1103C>G) (Table 1; Fig. S4, S5 & S6) and a novel in-frame deletion p.Val412del (c.1235_1237delTCG) in the highly conserved region within the *amrB* gene (Table 1; Fig. S4 & S7).

### amrB protein structure prediction by homology modelling and AlphaFold

The predicted homology modelled of the *B. pseudomallei* amrB protein found a homo-trimer protein that resembled the MexB protein, which is a part of the MexAB-OprM efflux pump in *P. aeruginosa* (Fig. 2A). It had a sequence identity score of 50.29%, a GMQE score of 0.77, a QSQE score of 0.79, and a QMEAN Z-score of -1.88. The Ramachandran plot showed that 94.1% of the residues were in the most favourable regions, with 1.66% of residues in the disallowed regions. These disallowed residues are mostly located in the loop region, which is commonly poorly defined. Most importantly, the amino acid threonine (Thr) is located in a Ramachandran-favoured area φ, ψ = (-63.33,-37.59) with confirmed α-helix chain configuration (Fig. S8), and hence it is considered reliable to be subjected to ΔΔG calculation in the mutant modelling analysis.

**Fig. 2.**
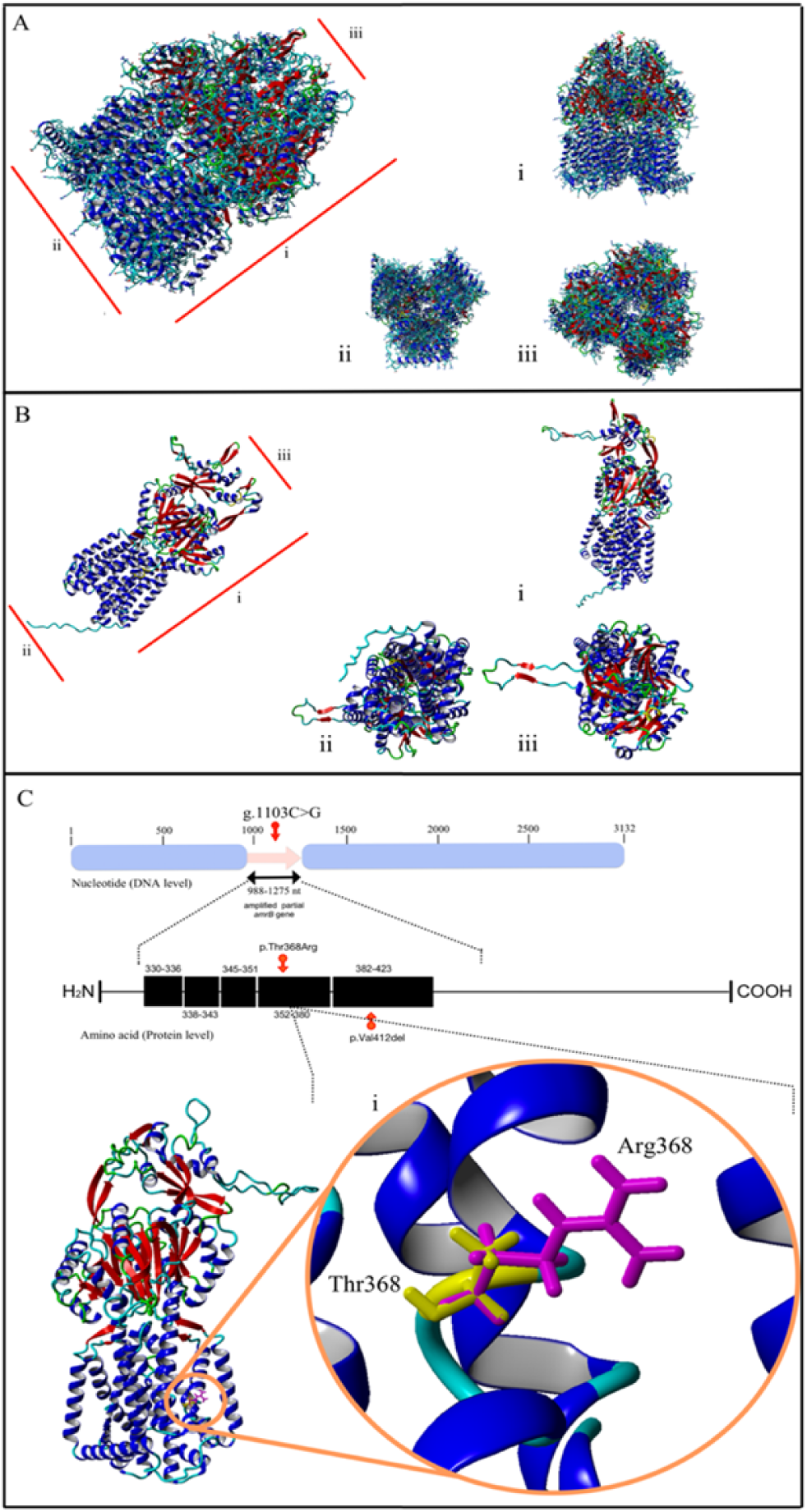
The homology-modelled predicted structure of the *B. pseudomallei* amrB protein, AlphaFold predicted structure of the *B. pseudomallei* amrB protein and the location of p.Thr368Arg, an amino acid substitution at the GEN^s^ *B. pseudomallei* amrB protein monomer. (A) Homology modelled structure of a homo-trimer amrB protein that resembles the *P. aeruginosa* multidrug exporter protein, mexB. i) side-view, ii) bottom-view, and iii) top-view. (B) AlphaFold predicted structure of a homo-trimer amrB protein that resembles the *P. aeruginosa* multidrug exporter protein, mexY. i) side-view, ii) bottom-view, and iii) top-view. (C) Ribbon diagram of p.Thr368Arg amino acid substitution in highly conserved region of *B. pseudomallei* amrB protein monomer. (i) A magnified view of the p.Thr368Arg mutation showing no amino acid side chain interaction perturbation. The side chain of Thr368 (GEN^r^), is shown in yellow, while side chain of Arg368 (GEN^s^) is shown in magenta. The highly conserved regions from amino acid 350 to 420 of the amrB protein are indicated as black blocks.

The structure of the amrB protein predicted by AlphaFold, on the other hand, is highly identical to the MexY protein, a subunit of the tripartite efflux pump protein, MexXY-OprM in *P. aeruginosa,* with a sequence identity of 70.2% (Fig. 2B). The protein sequence alignment E-value and bits score were 0.0 and 1453, respectively. Evaluation of the predicted protein exhibited a high confidence result with an overall pLDDT score of 89.6. The amino acid Thr at position 368 in the amrB protein sequence displayed a very high level of confidence with a pLDDT score of 92.48 (Fig. S9). Based on the PAE value scoring analysis, most of the residues in the amrB protein structure were classified as having a high level of confidence (green colour) particularly from the residues 300 to 400, including residue Thr368 (Fig. S10).

Despite the fact that the amino acid substitution p.Thr368Arg is located in a highly conserved region where any mutation, regardless of substitution, deletion, or addition of amino acid, can result in a malfunctional amrB protein, fascinatingly, we noted that the side chain of arginine (Arg) after substitution from Thr was observed not disrupting the protein-protein interaction or any van der Walls interaction, in contrast to the commonly found mutation in the highly conserved region, (Fig. 2C).

### Pairwise similarity alignment revealed that amrB protein structures predicted by homology modelling and AlphaFold are highly similar

Both protein structures solved by homology modelling and predicted by AlphaFold have been proven to possess similar structures using the superimposing method. This finding has increased our confidence in the reliability of both predicted structures to represent the real structure of the amrB protein, analogous to the structure solved experimentally with high reliability that can be used for *in silico* mutagenesis analysis.

TM-align pairwise similarity evaluation revealed that both of the amrB structures share a high similarity with a near perfect TM-score of 0.97639. The coverage of 98.1% (1025/1045) and the aligned residue fit within a low RMSD value (1.69 Å). It demonstrated that both structures possessed high geometry residue similarity with accurate structure superimposition (Fig. 3).

**Fig. 3.**
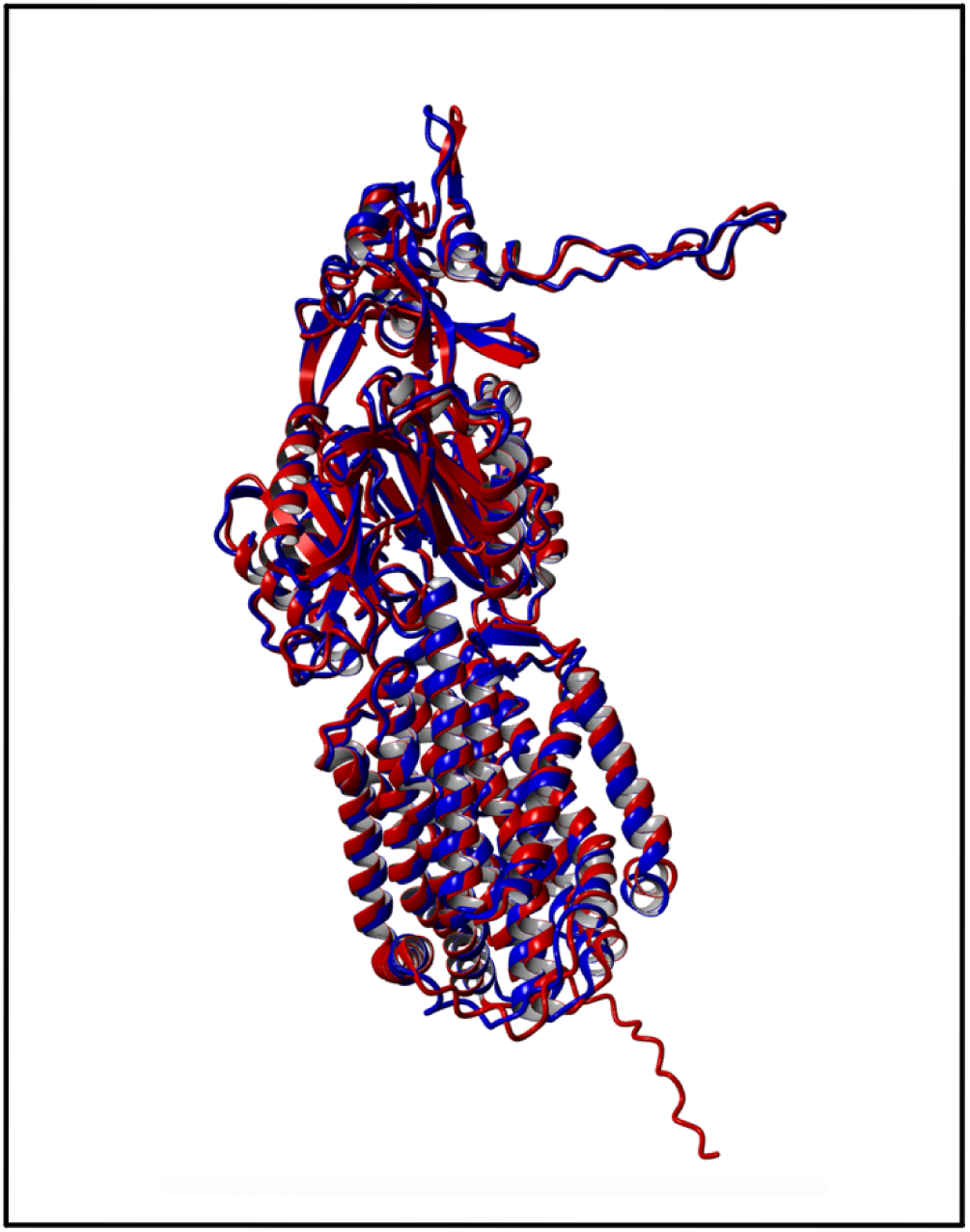
Superimposition of amrB protein domain structures predicted by the homology modelling and AlphaFold prediction showing high similarity with correct geometry. Superimposition result using TM-align (length of chain 1: 1045; length of chain 2: 1025; aligned length: 1025; RMSD: 1.69 Å; TM-score: 0.97639). Blue coloured structure was solved by homology modelling while red coloured structure was solved by the prediction using AlphaFold.

### The insignificant role of p.Thr368Arg in destabilising the amrB structure

The contribution of the novel in-frame deletion p.Val412del (c.1235_1237delTCG) in the highly conserved region to the production of a non-functional protein is regarded as clear and uncomplicated. Nevertheless, an intriguing discovery was made regarding the contribution of the amino acid substitution p.Thr368Arg to the function of the amrB protein. It has been demonstrated that the amino acid substitution p.Thr368Arg stabilises rather than destabilises amrB structures. Table 2 demonstrates that the missense mutation p.Thr368Arg stabilises both amrB protein structures predicted by homology modelling with a ΔΔG average value of -0.6 kcal/mol and by AlphaFold prediction with a ΔΔG average value of -1.2 kcal/mol. As depicted in Figure 2C, it may be because the Arg side chain is not involved in disrupting the protein-protein interaction. Therefore, we conclude that the amino acid substitution p.Thr368Arg technically has no effect on the structural stability perturbance of the amrB protein.

**TABLE 2.** The calculated ΔΔG caused by amino acid substitution p.Thr368Arg using homology modelled structure and AlphaFold predicted structure

| Run | Amino acid substitution | $\Delta\Delta G^a$<br>(kcal/mol) |
| --- | --- | --- |
| **Homology modelled structure** |  |  |
| 1 | p.Thr368Arg | -1.0 |
| 2 | p.Thr368Arg | -0.4 |
| 3 | p.Thr368Arg | -0.5 |
| 4 | p.Thr368Arg | -0.4 |
|  | Average | -0.6 |
| **AI AlphaFold predicted structure** |  |  |
| 1 | p.Thr368Arg | -1.0 |
| 2 | p.Thr368Arg | -1.3 |
| 3 | p.Thr368Arg | -1.2 |
| 4 | p.Thr368Arg | -1.1 |
|  | Average | -1.2 |

<sup>a</sup> $\Delta\Delta G$ calculated based on four protein models. $\Delta\Delta G=0$ corresponds to the same stability as the wild- type protein, $\Delta\Delta G \geq 0$ , the mutation is destabilising, $\Delta\Delta G \leq 0$ , the mutation is stabilising. A large ( $> 10$ kcal/mol) might indicate a problem in the overall structure of the model.

## DISCUSSION

To our knowledge, this is the first described observation of *in silico* mutational analysis of *B. pseudomallei* isolates that were unable to survive on the screening agar supplemented with GEN antibiotic and possessed low GEN MIC values, referred to as GEN^s^ *B. pseudomallei* strains, using homology-modelled and state-of-the art method, AlphaFold-solved amrB protein structures.

Apparently, there were no clinical symptoms or risk factors that lean toward either GEN^r^ or GEN^s^ strain infection, which indicates that a mutation at the efflux pump amrAB-OprA did not appear to reduce or increase GEN^s^ strain pathogenesis, analogous to the conclusions of previous studies (7, 9). Consequently, the application of differential diagnosis and screening agar supplemented with GEN antibiotic is insufficient to diagnose *B. pseudomallei* GEN^s^ strain infection due to the possibility of false-negative results, which justifies the importance of GEN^s^ prevalence determination in specific regions prior to the application of screening agar supplemented with GEN antibiotic to improve the accuracy of laboratory diagnosis. Intriguingly, the GEN^s^ *B. pseudomallei* strain has not yet been isolated in Sabah, despite the fact that it is a neighbouring state to Sarawak, corroborating previous research findings that GEN^s^ *B. pseudomallei* infection in Malaysian Borneo was restricted to Sarawak (6, 7). Future large-scale environmental and clinical screening studies would be crucial in confirming this hypothesis.

Hence, based on this finding, we concluded that the use of screening agar supplemented with GEN antibiotics in conjunction with symptoms and risk factors is still relevant in Sabah. However, it should be used with caution in regions where GEN^s^ strains are common, such as Sarawak, Malaysia, to avoid false-negative laboratory results and subsequent misdiagnosis of melioidosis disease as other bacterial infections or diseases with similar symptoms, such as *Pseudomonas sp.* infection (17), tuberculosis (68), or community-acquired pneumonia (16).

Based on the fact that a synonymous substitution within a gene can only reduce the rate of translation and increase the timing of protein folding (69, 70) and the location of the mutation outside the highly conserved region of the amrB protein, we affirmed that the c.1056T>G mutation reflects the *amrB* gene’s heterogeneity in *B. pseudomallei* and thus has no effect on the amrB protein’s function. On the contrary, although it appears most of small in-frame deletions or insertions have no or little effect on protein function, previous research has established the role of in-frame amino acid deletions in conferring a non-functional or partially functional protein (71–73). Furthermore, small in-frame deletions in selected genes have been found in patients with inherited eye disorders, such as childhood cataract and retinal dystrophy, indicating that in-frame deletion may play a role in phenotype abnormality (74). Recent finding showed that in-frame amino acid deletion played a role to cause milder muscular dystrophy as compared to the muscular dystrophy caused by the nonsense or frameshift mutation (75). These findings reinforced the potential role of pVal412del in contributing to a non-functional amrB protein and thus GEN susceptibility, in addition to the fact that this mutation is located in the highly conserved region of amrB protein.

In this study, the amrB structures predicted by homology modelling and AlphaFold were used instead of experimentally solved structures to calculate the ΔΔG value caused p.Thr368Arg amino acid substitution, similar to the previous studies method (76, 77) due to the lack of *B. pseudomallei* amrB protein crystal structures solved experimentally by X-ray crystallography, Nuclear Magnetic Resonance (NMR) spectroscopy, or electron microscopy methods that have been published in the Research Collaboratory for Structural Bioinformatics (RCSB) Protein Data Bank (PDB) to date [RCSB PDB database as of 24^th^ January 2023; Keywords: *B. pseudomallei* amrB, *B. pseudomallei* AmrAB-OprA and *B. pseudomallei* efflux pump]. The crystal structure solved by x-ray crystallography was the best choice for our homology modelling because it was recommended for calculating free energies with the FoldX tool (65) and for figuring out detailed mechanisms at the chemistry level with higher resolution (78).

The published guidelines for homology modelling recommend using structures with a resolution of higher than 2.6 Å as the template for mutagenesis analysis to ensure the high accuracy and reliability of the modelled protein’s atom locations (65). The MexB protein from *P. aeruginosa* was chosen as the template because it has the highest resolution crystal structure ever solved experimentally (2.7 Å) and shares 50.29% sequence identity with our amrB protein sequence. The achieved sequence identity was higher than the minimum requirement of 50% sequence identity with the template for providing a reliable protein homology model (47). Overall, the GMQE and QSQE scores indicated that the crystal structure of the amrB protein was accurate, with score values greater than 0.7. The QMQE score of 0.77 indicated that the homology-modelled protein’s tertiary structure was accurate (SWISS-MODEL, https://swissmodel.expasy.org/docs/help). On the other hand, a QSQE score of 0.79 showed that the prediction of the quaternary structure during the modelling process was accurate (SWISS-MODEL, https://swissmodel.expasy.org/docs/help) with a high degree of similarity between the template and the prediction in terms of stoichiometry and interfacial contacts (52).

Despite the fact that the QMEAN Z-score of -1.88 indicated that the predicted amrB protein structure was classified as average quality and slightly lower in quality than the experimental structures (54), the Ramachandran plot indicated that 94.17 percent of amino acids in amrB protein were located in the favoured residue region, which reflected that the predicted model was of high quality and had an adequate backbone geometry (56, 57). Furthermore, amino acid Thr at position 368 of the amrB protein sequence is located in the α-region at φ, ψ = (-63,-43), within the δ-region (delta/bridge region) in the Ramachandran plot, which was proposed to be a strongly favoured residue in real protein structure (55, 79), allowing for a reliable calculation of free energy change.

Similarly, the crystal structure of the amrB protein predicted using AlphaFold yielded a reliable result. Our predicted amrB protein structure received an overall pLDDT score of 89.6, indicating that the prediction was highly accurate with good backbone prediction and side chain orientation (80). Furthermore, amino acid Thr at position 368 of the amrB protein sequence, which has a pLDDT score of 92.48, indicating that this residue was predicted with high confidence (58), increased the reliability of ΔΔG values. The superimposition results showed that more than 98% of amino acid residues in both predicted crystal structures fit within 1.69 Å, implying that the proteins are nearly related and likely to perform the same biological function (58, 61, 81).

In this study, amino acid substitution p.Thr368Arg has been shown to stabilise amrB protein (homology modelling structure: -0.6 kcal/mol; AlphaFold structure: -1.2kcal/mol), in contrast to the previous conclusions that proposed amino acid substitutions, particularly in the highly conserved region with phenotypic changes are capable of causing massive destabilisation of protein structure (82–85). Recent published studies had successfully unravel the mystery of our finding, which concluded that although most of the amino acid substitutions with high ΔΔG, with ΔΔG value greater than or equal to 4.5 kcal/mol were almost always conferring to a non-functional protein (58% to 93%), while low ΔΔG (<4.5 kcal/mol) usually showed wild-type fitness activity, however, there were a number of substitutions with low ΔΔG, that did not significantly reduce protein stability but caused low fitness activity, conferring a non-functional protein through mechanisms other than stability and abundance reduction, such as mutation’s location at the active site residues in enzyme or at the key interaction site for binding or in the highly conserved region (25, 26).

Therefore, based on 1) our GEN-susceptibility phenotypic characterisation results, 2) the confirmed role of the amino acid substitution p.Thr368Arg in conferring GEN susceptibility by the previous experimental mutagenesis analysis (7), 3) amino acid substitution p.Thr368Arg is not located in the active site residue of an enzyme, 4) amino acid substitution p.Thr368Arg is not located at key interaction site for binding, 5) *in silico* analysis finding that confirmed the role of amino acid substitution p.Thr368Arg to stabilise the amrB protein structure and 6) amino acid substitution p.Thr368Arg is located in the highly conserved region of the amrB protein, we strongly proposed that amino acid substitution p.Thr368Arg plays a role in conferring GEN susceptibility through mechanisms other than stability perturbance, in which solely due to the location amino acid substitution in the highly conserved region. Besides that, we also suggest that amino acid substitution p.Thr368Arg might confer gentamicin susceptibility either singly or in combination with p.Val412del (c.1235_1237delTCG), an in-frame amino acid deletion in the highly conserved region of the amrB protein.

There were several limitations to this study, including the following: 1) The mutations identified in this study may not be the only ones discovered in GEN^s^ isolates, amrA and amrB genes are two main components in amrAB-OpA RND (20), in which mutation in amrA gene is still not yet explored, 2) Although the role of BpeAB-OprB efflux pump in extruding GEN antibiotic has been proven as strain-specific (21, 22), the mutation in this efflux pump that confer GEN-susceptibility using our clinical isolates has not yet to be determined. 3) The lack of an experimental crystal structure of amrB protein in the PDB forced the calculation of ΔΔG in this study to be based on the protein model predicted through homology modelling and AlphaFold. As a result, they may contain substantial errors. However, we verified that both structures were accurate prior to ΔΔG calculation using FoldX (86) and 4) The increased degree of freedom in the amino acid sidechain, involving amino acid Arg (position 362 in the amrB protein sequence), which is located close to the mutation site (position 368 in the amrB protein sequence), was expected to cause inconsistency in ΔΔG values between runs (FoldXYASARA, http://foldxyasara.switchlab.org/index.php/FoldX_tools_in_YASARA).

## CONCLUSION

Overall, 46 *B. pseudomallei* isolates from Malaysian Borneo (41 from Sabah and five from Sarawak) were successfully screened for GEN susceptibility, and three GEN^s^ isolates from Sarawak were phenotypically and genotypically characterised. In the highly conserved region, in-frame amino acid deletion p.Val412del and amino acid substitution p.Thr368Arg were found to confer GEN susceptibility. However, the role of the amino acid substitution p.Thr368Arg in conferring susceptibility was confirmed not by the disruption of the amrB protein’s overall structural stability, but rather by the mutation’s location in a highly conserved region. On the contrary, a synonymous mutation c.1056T>G in the less conserved region that translated to the similar amino acid, Ala, contributed to heterogeneity in the amrB gene of *B. pseudomallei* isolates.

## SUPPLEMENTAL MATERIAL

Supplemental material is available online only.

**SUPPLEMENTAL FILE 1**, PDF file, 16.7 MB.

## ACKNOWLEDGMENTS

The authors would like to thank the Director General of Health Malaysia for the permission to publish this article. We would also like to thank Dr. William Gotulis, Director of Queen Elizabeth Hospital (QEH), Dr. Ahmad Toha Samsudin and all Microbiology Unit Staff, Department of Pathology, Queen Elizabeth Hospital, for their unwavering technical support of this research.

This research received no specific grant from any funding agency in the public, commercial, or not-for-profit sectors.

A.K. contributed to the study conceptualisation, formal analysis, investigation, methodology, software, validation, visualization, writing-original draft and writing-review & editing. S.N. contributed to the conceptualization, funding acquisition, investigation, project administration, resources, supervision and writing-review & editing. M.A.S contributed to the conceptualization, funding acquisition, investigation, project administration, resources and writing-review & editing. M.Y.N.R and M.Y.Z. participated in investigation, methodology, writing-original draft and writing-review & editing. N.A.N.K participated in the investigation and writing-review & editing. NI contributed to the conceptualization, funding acquisition, investigation, methodology, project administration, resources, supervision, validation, visualization and writing-review & editing. All authors were involved in counterchecking and approving the final version of the manuscript.

We are not aware of any affiliations, memberships, funding, or financial holdings that might affect the objectivity of this article. We declare that there is no conflict of interest regarding the publication of this article.

